# Yes-associated Protein Induces Age-dependent Inflammatory Signaling in the Pulmonary Endothelium

**DOI:** 10.1101/2025.02.26.640349

**Authors:** Memet T. Emin, Alexandra M. Dubuisson, Prisha Sujin Kumar, Carsten Knutsen, Cristina M. Alvira, Rebecca F. Hough

**Affiliations:** Pediatric Critical Care, Hospitalist, and Palliative Medicine, Department of Pediatrics, Columbia University Irving Medical Center; Louisiana State University Health Sciences Center Shreveport; Department of Pediatrics, University of California, San Francisco School of Medicine

**Keywords:** Acute Lung Injury, PARDS, lung endothelium, Yes-associated protein, NF-κB

## Abstract

Acute Lung Injury (ALI) causes the highly lethal Acute Respiratory Distress Syndrome (ARDS) in children and adults, for which therapy is lacking. Children with Pediatric ARDS (PARDS) have a mortality rate that is about half of adults with ARDS. Improved ALI measures can be reproduced in rodent models with juvenile animals, suggesting that physiologic differences may underlie these outcomes. Here, we show that pneumonia-induced ALI caused inflammatory signaling in the endothelium of adult mice which depended on Yes-associated protein (YAP). This signaling was not present in 21-day-old weanling mice. Transcriptomic analysis of lung endothelial responses revealed nuclear factor kappa-B (NF-*κ*B) as significantly increased with ALI in adult versus weanling mice. Blockade of YAP signaling protected against inflammatory response, hypoxemia, and NF-*κ*B nuclear translocation in response to *Pseudomonas aeruginosa* pneumonia in adult mice. Our results demonstrate an important signaling cascade in the lung endothelium of adult mice that is not present in weanlings. We suggest other pathways may also exhibit age-dependent signaling, which would have important implications for ARDS therapeutics in the adult and pediatric age groups.

## Introduction

Acute lung injury (ALI) causes the Acute Respiratory Distress Syndrome (ARDS) and Pediatric ARDS (PARDS). Mortality rates in adult patients with ARDS are double those of PARDS patients (1–3). Preclinical studies in rodents show decreased lung injury measures in juvenile mice (4–6), suggesting that age-specific physiological differences may underlie the distinct outcomes in ARDS and PARDS. Understanding the mechanisms of this differences may aid in the development of therapeutics for both age groups.

The Hippo pathway and its main effectors Yes-associated Protein (YAP) and PDZ-binding motif (TAZ) are vital for growth and differentiation (7–9). YAP is essential for the development of the endothelial barrier in the retina and brain (10), and interacts with tight junction components in the lung epithelium (11). Recently, YAP was shown to protect against lung injury (12, 13). Since YAP is developmentally important and implicated in ALI, we hypothesized that YAP responses to ALI could differ in juvenile and adult rodents.

Here, we show robust YAP protein expression in response to *Pseudomonas aeruginosa* infection in adult mice, but not in weanlings. YAP knockdown prevented the inflammatory response and hypoxemia in adult mice, suggesting a causal role of YAP in our injury model. Knockdown of YAP decreased nuclear translocation of NF-*κ*B in adult mice, suggesting that YAP contributes to pro-inflammatory signaling in the pulmonary endothelium of adult but not juvenile mice.

## Methods

### Materials

For immunoblot, YAP mouse antibody (Cat#12395, 1:1000), YAP rabbit antibody (cat#14074, 1:1000), GAPDH mouse antibody (Cat#97166, 1:1000), and NF-*κ*B p65 rabbit antibody (Cat#8242, 1:1000) were purchased from Cell Signaling Technology; HDAC1 mouse monoclonal antibody (Cat# 66085, 1:1000) was purchased from Proteintech; and actin rabbit antibody (cat# A2066, 1:1000) was purchased from Sigma-Aldrich. For lung cell isolation, CD31 rat monoclonal antibody clone MEC 13.3 (1:500, catalog 557355) was purchased from BD Biosciences. CD45 rat monoclonal antibody (1:600, catalog 14-0451-85), CD41 rat monoclonal antibody (1:500, catalog 14-0411-85), and CD326 rat monoclonal antibody (1:500, catalog 14-5791-85) were purchased from Invitrogen. siRNA construct targeting Yap1 (siRNA ID #s76160) was purchased from ThermoFisher Scientific. Male and female C57BL/6J mice were purchased from Jackson Laboratory (RRID:IMSR_JAX:000664). *P. aeruginosa* strain K was obtained from Alice Prince (Columbia University). All protocols were reviewed and approved by the Columbia University Institutional Animal Care and Use Committee.

### Pseudomonas-induced lung injury

*P. aeruginosa* was prepared as previously described (14). We anesthetized mice and intranasally (i.n.) instilled 2.5 × 10^5^ colony forming units (CFUs) *P. aeruginosa* at exponential growth phase in 50µl sterile PBS (adult) or 20µl (weanling). 24 hours later, we anesthetized the mice, performed a tracheotomy, and collected BAL as described (14). We retained lungs after BAL for downstream experiments. Extra-vascular lung water (EVLW) was measured as described (14, 15).

### SpO_2_ measurement

For non-invasive O_2_ saturation (SpO_2_) measurements, we anesthetized mice and removed the fur from right thigh. We placed a MouseOx® Pulse Oximeter sensor (Starr Life Sciences Corp, Oakmont, PA) on the right femoral artery region and recorded SpO_2_ after a 15-min stabilization period.

### YAP RT-PCR

We isolated RNA from whole lung using miRNeasy kit (Qiagen) according to the manufacturer’s protocol. 250ng of RNA was reverse-transcribed into cDNA. We performed quantitative PCR using TaqMan Gene Expression Master Mix with QuantStudio™ 5 Real-Time PCR Instrument (Thermo Fisher Scientific Applied Biosystems). The following primers were used: Yap1: Assay ID Mm01143263_m1 and Actb: Assay ID Mm02619580_g1 (ThermoFisher).

### YAP knockdown

Anesthetized mice were in vivo transfected with YAP siRNA (50*μ*g in 150*μ*L) by tail vein injection 24 hours before *P. aeruginosa* instillation.

### Immunoblot

Without freezing samples, we enriched lungs for cytoplasmic and nuclear fractions using NE-PER Nuclear and Cytoplasmic Extraction Reagents (Thermo Fisher Scientific, cat #78833). Equal amounts of protein were separated by SDS-PAGE, transferred onto nitrocellulose membrane, blocked in StartingBlock Blocking Buffer (Thermo Scientific / Pierce, catalog 37543), and immunoblotting was performed.

### Endothelial cell isolation

Endothelial cells from C57BL/6J mice were isolated as described (16, 17) with the following modifications. Lungs were digested for 15 min at 37°C in 0.38mg/ml Liberase (Roche) in D-PBS + 0.1% albumin. We used a DynaMag-2 Magnet and Dynabeads Sheep Anti-Rat IgG (Invitrogen) to deplete CD45+ and CD41+ cells, then positively enrich three times for CD31+ cells. These procedures yielded ∼10^5^ CD31+ cells per mouse lung with <5% leukocyte contamination.

### Bulk RNA-sequencing

RNA was isolated from freshly isolated endothelial cells bound to magnetic beads (miRNeasy). RNA-sequencing was performed by Azenta. We aligned sequences to the GRCm39 mouse genome using STAR aligner (v2.10.a) (18). We analyzed differential gene expression (DE) with EdgeR (3.38.4) (19) using a generalized linear model with thresholds for significance set at <0.05 FDR (Benjamani-Hochenberg) and 1>|Log2(FC)|. Pathway analysis was performed using Metascape (v3.5) (20), using up- and down-regulated genes separately. Code for this analysis can be found at https://github.com/CarstenKnutsen/bulkrnaseq_pseudomonas_lung_ec/.

## Results

### Differential lung injury responses in weanling and adult mice

To determine lung injury responses, we intranasally instilled weanling (P21-24) and adult (8-10 week old) mice with *P. aeruginosa*. Twenty-four hours later, we assessed mice for lung injury by measurement of BAL protein, BAL cell count, BAL neutrophils, EVLW, BAL cytokines, and non-invasive measurement of SpO_2_. Following *P. aeruginosa* instillation, adult mice had an increase in BAL protein, BAL cell count, BAL neutrophils, EVLW, BAL Il-6 and BAL TNF-*α* (**Fig 1A-E**), and a decrease in SpO_2_ (**Fig 1F**) compared to weanling mice. Saline treatment did not produce different responses in any lung injury measurements.

**Figure 1.**
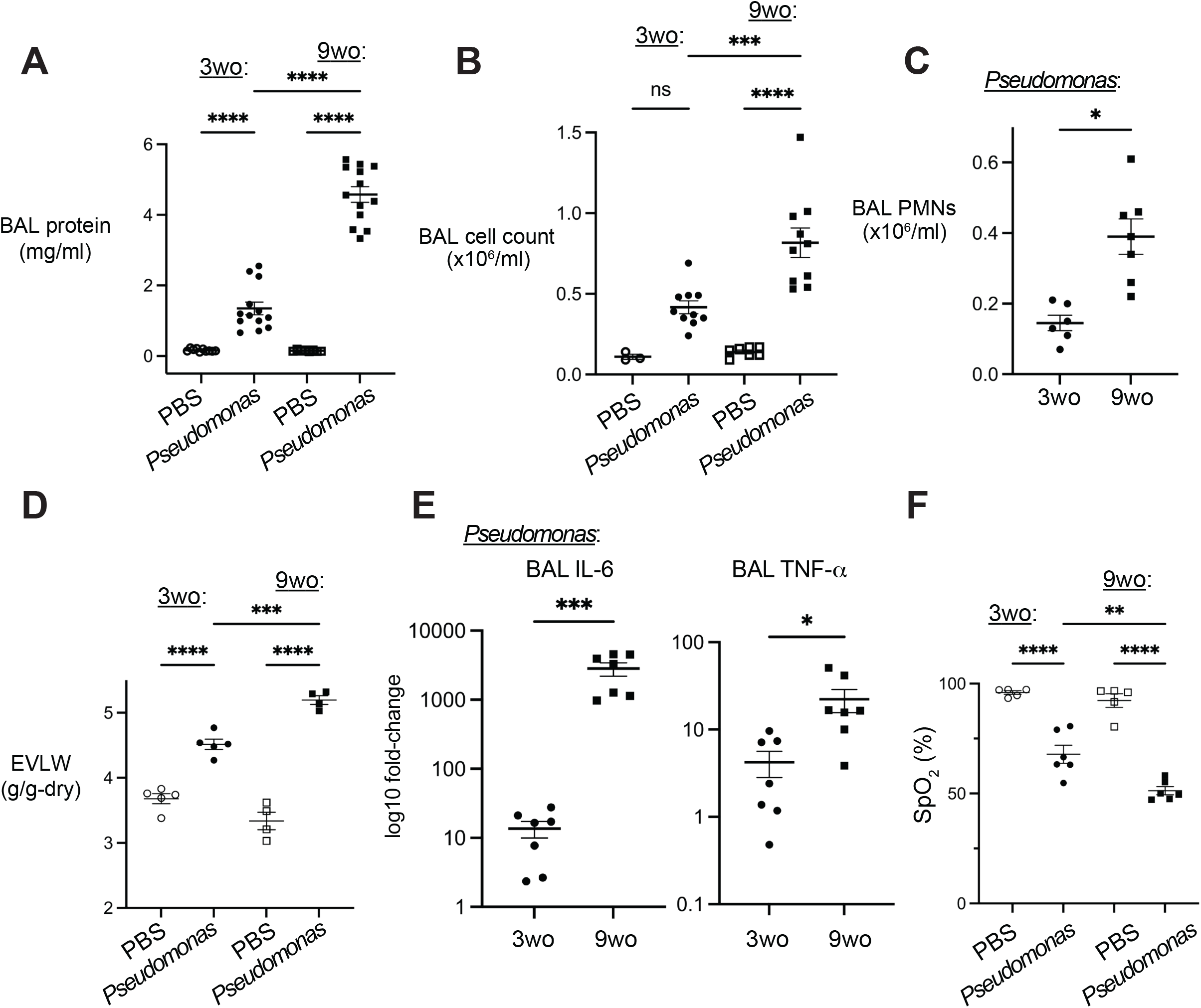
Responses to intranasal *Pseudomonas aeruginosa* in weanling and adult mice. Analyses were performed 24h after intranasal instillation of 2.5 × 10^5^ CFU of *P. aeruginosa* or PBS. (**A-C, E**) BAL was obtained by instillation of ice cold PBS intratracheally. (**A**) BAL protein was determined by BCA assay. (**B-C**) BAL cell and neutrophil counts were determined with Vetscan hematology analyzer. (**D**) ELVW was determined in separate experiments. (**E**) BAL cytokines were measured by multiplex array (Eve Technologies). (**F**) SpO_2_ was determined in anesthetized mice using a MouseOx® Pulse Oximeter. Data are shown as mean±SE. Number of replicates are indicated by dots (≥4 for all groups). *BAL*, bronchoalveolar lavage; *3wo*, 3 week-old mice; *Pseudomonas, Pseudomonas aeruginosa* strain K; *PMNs*, neutrophils; *EVLW*, extravascular lung water; *IL-6*, interleukin-6; *TNF-α*, tumor necrosis factor *α*; *SpO*_*2*_, non-invasive O_2_ saturation. \**p*<0.05, \*\**p*<0.01, \*\*\**p*<0.001, \*\*\*\**p*<0.0001. Multiple comparisons were analyzed by two-way ANOVA; paired differences were compared by unpaired 2-tailed t test.

Given the difference in size of the weanling and adult mice (∼10g vs. ∼25g), we considered whether delivery of *P. aeruginosa* could differ between the two age groups. We also assessed for altered clearance of *P. aeruginosa*, as decreased clearance of *P. aeruginosa* has been noted in mice <20 days old (21). To assess *P. aeruginosa* delivery, we instilled the same number of CFUs in the same volume to both age groups, and again in two different volumes. The lung injury results did not differ when the volume was changed. To assess bacterial clearance, we measured BAL CFUs 24h after instillation and noted no difference between the two age groups (**Supplemental Fig 1-https://figshare.com/s/5e58428fdf883f37fe84**). These findings suggest that a similar number of CFUs was delivered in both ages and that bacterial clearance did not differ between the age groups.

### P. aeruginosa-induced changes in YAP protein expression differ with age

We determined changes in expression of YAP in response to *P. aeruginosa*-induced lung injury. Following BAL and a vascular wash, we enriched whole lungs from *P. aeruginosa*- and saline-treated mice into nuclear and cytoplasmic portions. We found that adult mice had robust increases in YAP protein expression in response to *P. aeruginosa* instillation in both cytoplasmic and nuclear fractions (**Fig 2A, B**). In contrast, weanling mice had decreased cytoplasmic expression of YAP and no change in nuclear expression. Of note, there was no difference in YAP protein expression between age groups in saline-treated controls (**Supplemental Fig 2**), suggesting that baseline YAP expression does not differ between the two age groups. We assessed YAP mRNA expression with RT-PCR and noted that YAP mRNA expression in both weanling and adult mice decreased upon *P. aeruginosa*-induced injury (**Supplemental Fig 3**), suggesting a post-transcriptional mechanism of increased YAP protein expression in adult mice.

**Figure 2.**
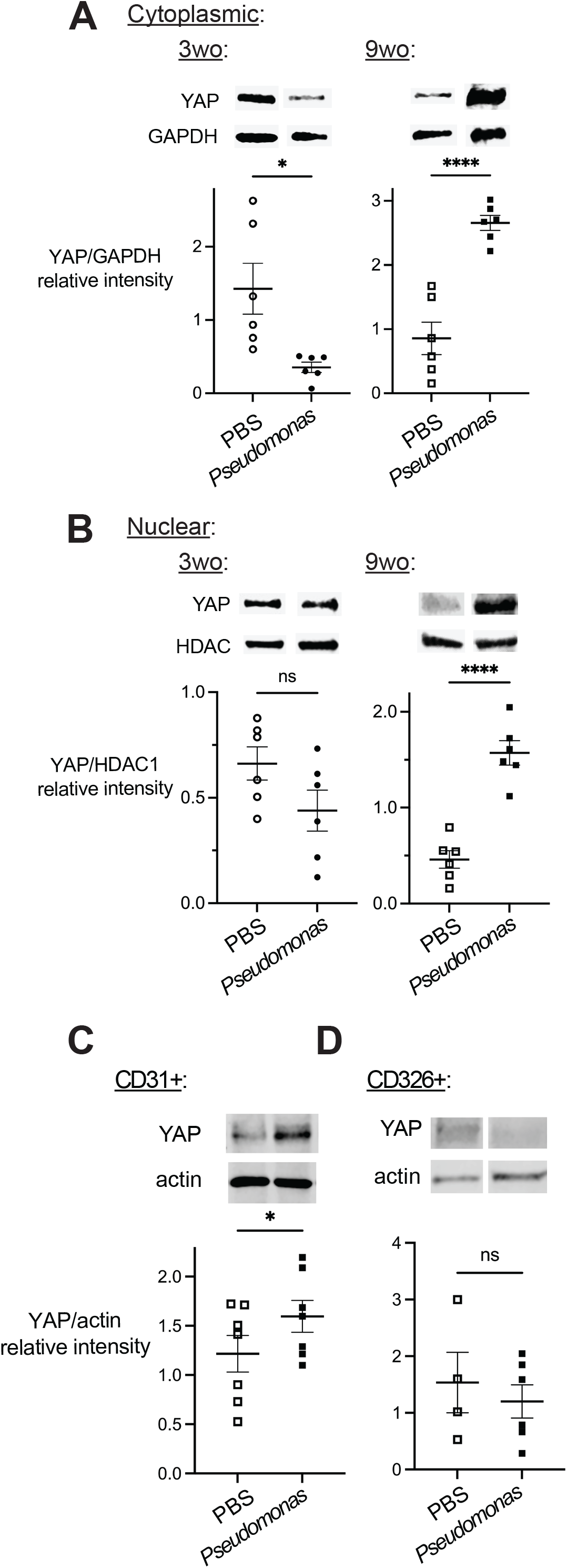
YAP expression responses to *P. aeruginosa*-induced lung injury in weanling and adult mice. Representative gels and scatter plots show immunoblotting and densitometry of enriched cytoplasmic (**A**) and nuclear (**B**) fractions of whole lung, and enriched lung endothelial cells (**C**) and epithelial cells (**D**). All animals were treated with i.n. PBS or *P. aeruginosa* 24 hours before lung removal. Antibodies used were YAP rabbit monoclonal, GAPDH mouse monoclonal, HDAC mouse monoclonal, and actin rabbit polyclonal. Lanes were run on the same gel. Number of replicates are indicated by dots (≥4 for all groups). *3wo*, 3 week-old mice; *n*.*s*., not significant. \**p* < 0.05, \*\*\*\**p*<0.0001. Differences between groups were compared by t test.

### P. aeruginosa-induced YAP protein increases occur in lung endothelial cells

Given the substantial increase in inflammatory cells in the BAL of adult versus weanling mice (**Fig 1B-C**), we considered whether leukocyte expression of YAP (22) contributed to YAP increases in adult mice. To determine the cell type in which YAP expression increased, we enriched lungs for three fractions: endothelial cells, epithelial cells, and leukocytes plus platelets. We performed immunoblot for YAP in each of the enriched cell populations. We noted an increase in YAP protein expression in endothelial cells (**Fig 2C**). There was no significant increase in YAP expression in epithelial cells (**Fig 2D**), and no YAP protein expression was detected in the leukocyte- and platelet-enriched fraction in PBS or *P. aeruginosa*-treated mice. These data suggest that YAP increases in lung endothelial cells are the main driver of the increased YAP in adult mice.

### Vascular YAP mediates P. aeruginosa-induced lung injury in adult mice

Since YAP expression increases in adult mice, but not in weanling mice, we hypothesized that adult mice exhibit YAP-dependent lung injury responses that weanling mice lack. Thus, we targeted endothelial YAP in adult mice by tail vein injection of siRNA-targeted YAP. Injection of YAP-targeted siRNA decreased YAP expression in *P. aeruginosa* infected mice by ∼80% (**Fig 3A**). We assessed lung injury measures as in **Fig 1** and found that YAP knockdown prevented *P. aeruginosa*-induced increases in BAL cell count and IL-6, as well as *P. aeruginosa*-induced decreases in SpO_2_ (**Fig 3B-D**). This suggests that endothelial YAP expression is important for inflammatory and physiological responses in *P. aeruginosa*-induced lung injury. Notably, YAP knockdown had no effect on BAL protein and EVLW, suggesting that endothelial YAP may not be essential for barrier failure.

**Figure 3.**
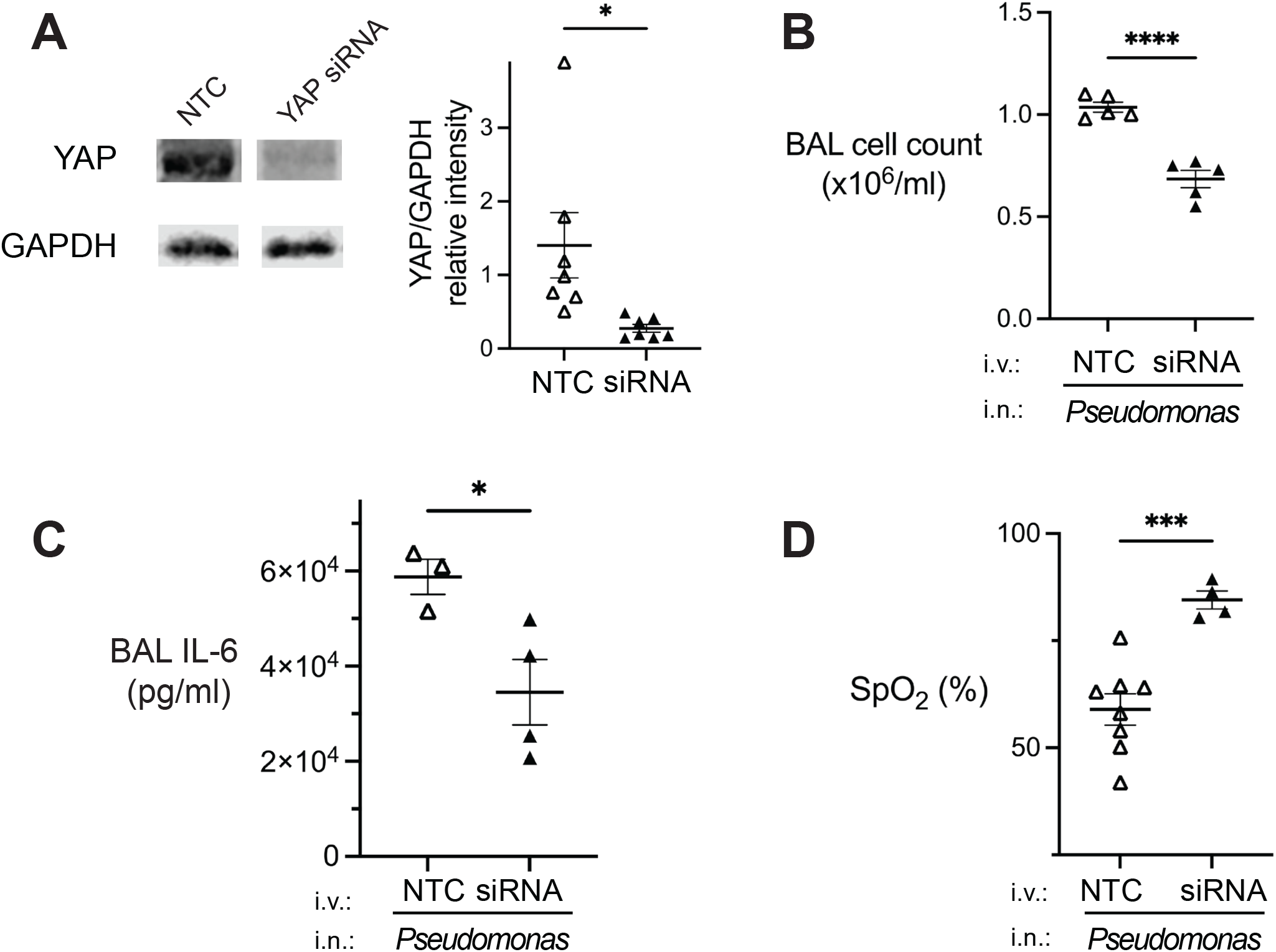
YAP mediates *P. aeruginosa*-induced lung injury in adult mice. Mice were injected intravenously with YAP-targeted siRNA or non-targeting control siRNA (NTC) 24h before all mice were treated with *P. aeruginosa* intranasal instillation. (**A**) Representative gel and scatter plots show immunoblotting and densitometry of whole lung cytoplasmic fraction. Antibodies used were YAP rabbit monoclonal and GAPDH mouse monoclonal. There was high variability of lung YAP expression, but statistical significance remained when the highest outlier from the NTC-treated group was removed from the analysis. (**B**) BAL cell counts were determined with Vetscan hematology analyzer. (**C**) BAL IL-6 was measured by multiplex array. (**D**) SpO_2_ was determined using a MouseOx® Pulse Oximeter. Number of replicates are indicated by triangles (≥3 for all groups). *BAL*, bronchoalveolar lavage; *3wo*, 3 week-old mice; *IL-6*, interleukin-6; *TNF-α*, tumor necrosis factor *α*; *SpO*_*2*_, non-invasive O_2_ saturation. \**p*<0.05, \*\*\**p*<0.001, \*\*\*\**p*<0.0001. Paired differences were compared by t test.

### YAP expression drives endothelial NF-κB responses in P. aeruginosa-infected adult mice

To determine differences in *P. aeruginosa*-induced lung injury endothelial responses between weanling and adult mice, we performed bulk RNA sequencing on freshly isolated endothelial cells from mice treated with i.n. *P. aeruginosa* and control mice treated with PBS. Gene expression differences between both age groups did not differ greatly under control conditions (107 genes upregulated; 128 downregulated, **Fig 4A, Supplemental data files, Gene Expression Omnibus GSE291989**). However, gene expression responses to *Pseudomonas*-induced lung injury differed greatly between the age groups (966 upregulated; 1474 downregulated). This suggests substantial differences in lung endothelial responses to inflammatory injury of weanling and adult mice.

**Figure 4.**
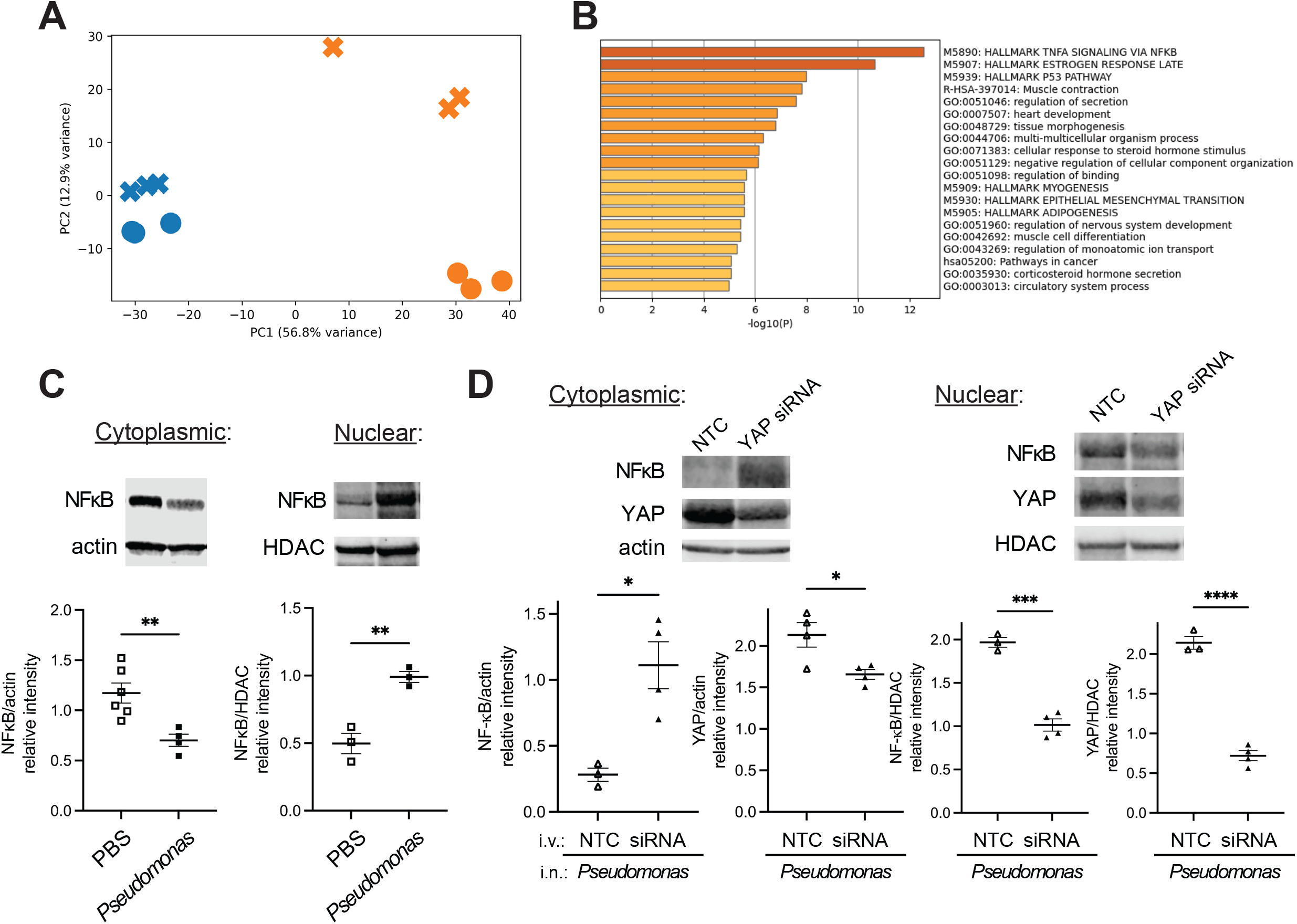
YAP expression drives endothelial NF-*κ*B responses in *P. aeruginosa*-infected adult mice. (**A**) Principal component analysis of four treatment groups. *PC1*, principal component 1; *blue*, saline-treated; *orange, P. aeruginosa*-treated. •, adult; **x**, weanling. (**B**) Top pathways in which adult response to *P. aeruginosa* instillation is higher than in weanling mice. (**C**) Representative gel and scatter plots show immunoblotting and densitometry of whole lung cytoplasmic and nuclear enriched fractions: (**C**) 24h after PBS or *P. aeruginosa* instillation and (**D**) following pretreatment with YAP siRNA by tail vein injection and *P. aeruginosa* instillation. Antibodies used were NF-*κ*B p65 rabbit monoclonal, YAP rabbit monoclonal, HDAC mouse monoclonal, and actin rabbit polyclonal. Number of replicates are indicated by dots (≥3 for all groups). *NF-κB*, nuclear factor-kappa B; *NTC*, non-targeting control siRNA; *siRNA*, YAP-targeted siRNA. \**p*<0.05, \*\**p*<0.01, \*\*\**p*<0.001, \*\*\*\**p*<0.0001. Paired differences were compared by t test.

Of note, there was no significant alteration in *Yap1* gene expression with *P. aeruginosa* instillation in either weanling or adult endothelia, consistent with our RT-PCR results (**Supplemental Fig 3**). We identified pathways that exhibited increased expression in adult mice in response to *P. aeruginosa*-induced lung injury relative to weanling mice. The most significant increases were found in TNF*α* signaling via NF-*κ*B (specific genes: *Klf9, Cdkn1a, Map3k8, Rcan1, Ets2, Hes1, Gadd45b, Nfkbia, Nr4a2, Per1, Plaur, Tgif1, Socs3, Ccrl2, Plk2, Kdm6b, Ppp1r15a, Maff, Ccnl1, Zbtb10, Sik1*, **Fig 4B, Supplemental data files**). Although both weanling and adult endothelial cells increased their expression of genes involved in this pathway, the adult endothelial responses were increased relative to responses in weanling mice.

We considered whether YAP drove NF-*κ*B nuclear translocation in endothelia, leading to a more robust inflammatory response in adult mice. First, we determined whether *P. aeruginosa* instillation drove NF-*κ*B translocation to the nucleus in adult mice. 24h after *P. aeruginosa* instillation, we enriched whole lung into nuclear and cytoplasmic fractions. We noted a significant increase in NF-*κ*B in the nuclear fraction and a decrease in the cytoplasmic fraction (**Fig 4C**). In weanlings, there was an increase in NF-*κ*B in the nuclear fraction, but no significant difference in NF-*κ*B quantity in the cytoplasmic fraction upon instillation of *P. aeruginosa* (**Supplemental Fig 4**), suggesting that NF-*κ*B nuclear translocation in response to *P. aeruginosa* in weanlings may be decreased in comparison to adult mice. To determine whether the increase in YAP expression in adult mice drove NF-*κ*B translocation, we knocked down YAP expression by vascular injection of siRNA as in **Fig 3A** and instilled *P. aeruginosa* into control- and siRNA-treated mice 24h later. 24 hours after *P. aeruginosa* instillation, we measured NF-*κ*B in the nuclear and cytoplasmic fractions. We noted that YAP knockdown increased NF-*κ*B protein expression in the cytoplasm and decreased NF-*κ*B expression in the nucleus (**Fig 4D**), suggesting that YAP mediated NF-*κ*B nuclear translocation.

## Discussion

Although pneumonia is a common cause of ARDS and PARDS, ALI signaling including barrier failure and infiltration of inflammatory cells that ultimately leads to hypoxemia is orchestrated by the microvascular endothelium (15, 23). Here, we present a novel mechanism of age-dependent signaling in the pulmonary endothelium. Our data suggest that in adult mice YAP-dependent inflammatory signaling contributes to ALI, worsening outcomes. One possibility is that bacterial clearance of *P. aeruginosa* differs in the two age groups. Indeed, decreased clearance of *P. aeruginosa* has been noted in mice >20 days old (21). While we did see decreased neutrophilic infiltration in weanling mice (**Fig 1C**), there was no significant difference in BAL colony forming units at 24h (**Supplemental Fig 1**), suggesting that bacterial clearance did not differ significantly between the age groups.

Previously work using models of indirect lung injury showed that endothelial YAP protected against lung injury. Indeed, YAP inhibited NF-*κ*B nuclear translocation by promoting degradation of tumor necrosis factor receptor-associated factor 6 (TRAF-6) in these models (13). In the epithelium, nuclear YAP may increase *ikba* gene expression, also inhibiting NF-*κ*B nuclear translocation (24). Endothelial YAP was also recently shown to protect against lung injury secondary high-tidal volume ventilation (12). In our transcriptomics dataset, expression of the *ikba* mouse homologue *nfkbia* was induced in both weanling and adult mice, supporting YAP-independent drivers of transcription. It is well-known that YAP has a pleiotropic role in lung injury and repair. We suggest that knockdown, rather than knockout of YAP used in these prior studies, more accurately reflects a possible pharmacological inhibition of YAP that could be used in pre-clinical and/or clinical trials. Furthermore, complete inhibition of YAP raises concerns about the effects on development, differentiation, and repair that are attributed to YAP. Indeed, it is possible that YAP knockout leads to up-regulation of alternative pathways (25). Our work suggests that the role of YAP in lung injury and repair is complex and requires further study.

In neonatal mice in the early alveolar stage of lung development, NF-*κ*B is known to play a protective role, ultimately promoting angiogenesis (17, 26–28). To our knowledge, this role has not been described in weanling mice in the late alveolar stage used in our study. However, it is possible that NF-*κ*B nuclear translocation in weanling mice results in signaling that differs from canonical pro-inflammatory signaling in adults.

Our study has limitations. First, the signaling that leads to differential YAP expression between the weanling and adult age groups is unclear. Our RT-PCR (**Supplemental Fig 3**) and transcriptomics data suggest that the regulation is post-transcriptional, yet the drivers of these differences remain unknown. Second, our bulk RNA sequencing data do not reveal the specific endothelial cell types that drive the inflammatory responses. We and others have identified the post-capillary venule as a primary location of inflammatory signaling (29–33, 14). However, a gene expression signature of this cell type to our knowledge remains elusive. Finally, the mechanism(s) by which YAP drives NF-*κ*B nuclear translocation in adult mice remains unclear. This deserves further investigation.

Here, we show evidence for novel age-dependent inflammatory signaling in the pulmonary endothelium. Our work suggests that targeting of YAP-dependent endothelial signaling may be helpful in the prevention and treatment of ARDS but may be detrimental in PARDS. Our transcriptomics dataset suggests numerous differences in inflammatory responses in the pulmonary endothelium between weanling and adult age groups, of which YAP-driven NF-*κ*B is but one example. Further investigation into age-dependent mechanisms of lung injury could aid in the tailoring of therapies across the lifespan.

## Acknowledgements

We are grateful to Jahar Bhattacharya, Wellington V. Cardoso, Mohammad N. Islam, and Galina Gusarova for contributive discussions.

