## Supplemental Figures for "Yes-associated Protein Induces Age-dependent Inflammatory Signaling in the Pulmonary Endothelium"

**Supplemental Figure Legends**

**Supplemental Figure 1. *Pseudomonas aeruginosa* in BAL of weanling and adult mice.** 24h after instillation of *P. aeruginosa* (2.5 x $10^5$ CFU), BAL was collected from anesthetized mice by intratracheal instillation of 0.7ml of ice cold D-PBS. BAL was cultured overnight on agar plates without antibiotics. *n.s.*, not significant. Data are shown as mean ± SEM. n=7. Differences between groups were compared by t test.

**Supplemental Figure 2. Expression of YAP in weanling and adult mice treated with PBS.** Representative gels and scatter plots show immunoblotting and corresponding densitometry of enriched cytoplasmic and nuclear fractions of whole lung. All animals were treated with i.n. PBS 24 hours before lung removal. Antibodies used were YAP rabbit monoclonal, GAPDH mouse monoclonal, and HDAC mouse monoclonal. Lanes were run on the same gel. *3wo*, 3 week-old mice; *n.s.*, not significant. Data are shown as mean ± SEM. n≥4 as indicated by dots. Differences between groups were compared by t test.

**Supplemental Figure 3. mRNA expression of Yes-associated protein in lung of weanling and adult mice.** 24h after intranasal instillation of 2.5 x $10^5$ CFU of *P. aeruginosa* or PBS as indicated, anesthetized mice were subjected to BAL and vascular washing with ice-cold PBS. Lungs were collected for RNA extraction (Qiagen) and YAP RT-PCR was performed. Data are shown as mean±SE. n≥4 for all groups. *BAL*, bronchoalveolar lavage; *3wo*, 3 week-old mice; *Pseudomonas*, *Pseudomonas aeruginosa*. ***p=0.0005, ****<0.0001. Paired differences were compared by t test.

**Figure 4. Endothelial NF-κB responses in *P. aeruginosa*-infected weanling mice.** Mice were treated with intranasal instillation of 2.5 x $10^5$ CFU of *P. aeruginosa* or PBS as indicated. 24h later, BAL was removed, vasculature was washed, and lung was enriched for cytoplasmic and nuclear fractions. Representative gel and scatter plots show immunoblotting and densitometry. Antibodies used were NF-κB p65 rabbit monoclonal, HDAC mouse monoclonal, and actin rabbit polyclonal. Data are shown as mean±SE. Number of replicates are shown by circles (≥3 for all groups). *Pseudomonas*, *Pseudomonas aeruginosa; NF-κB, nuclear factor-kappa B;* **$p$<0.01; *ns*, not significant. Paired differences were compared by t test.

**Supplemental Figure 1:** *Pseudomonas aeruginosa* in BAL of weanling and adult mice

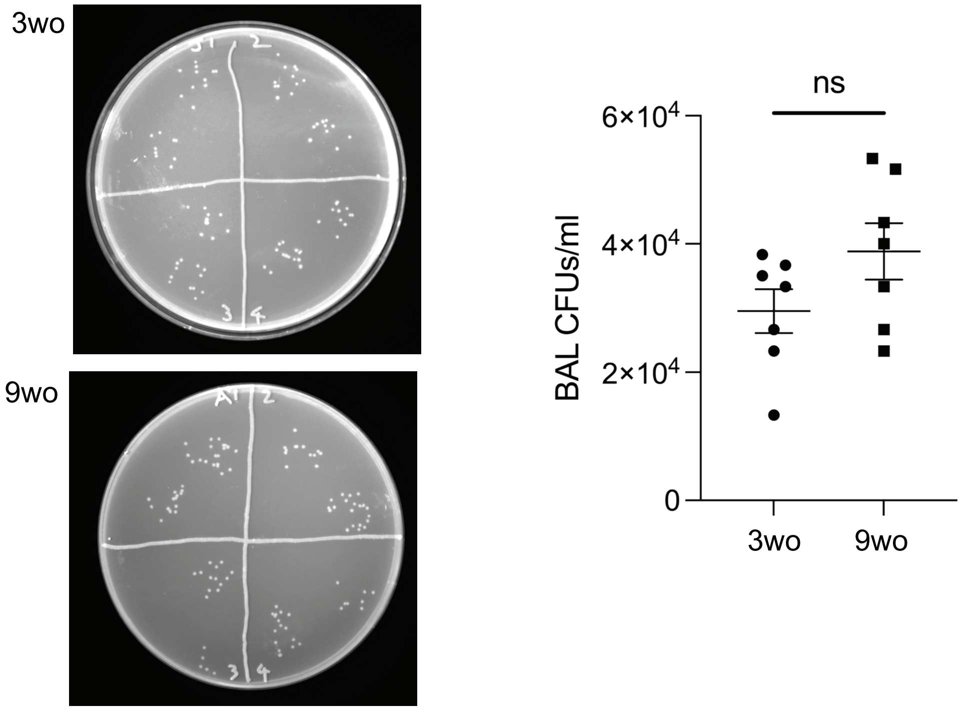

**Supplemental Figure 2:** Expression of YAP in weanling and adult mice treated with PBS

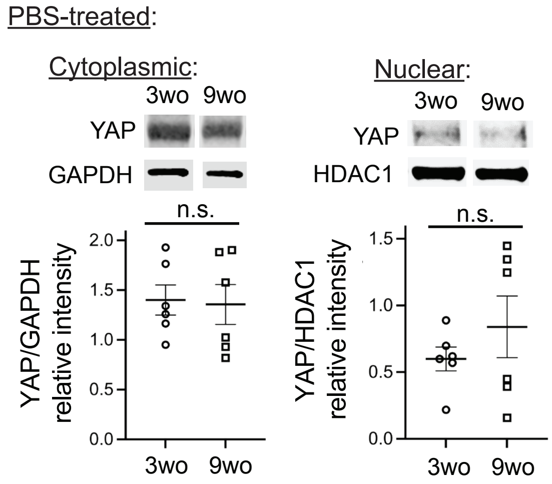

**Supplemental Figure 3:** mRNA expression of Yes-associated protein in lung of weanling and adult mice

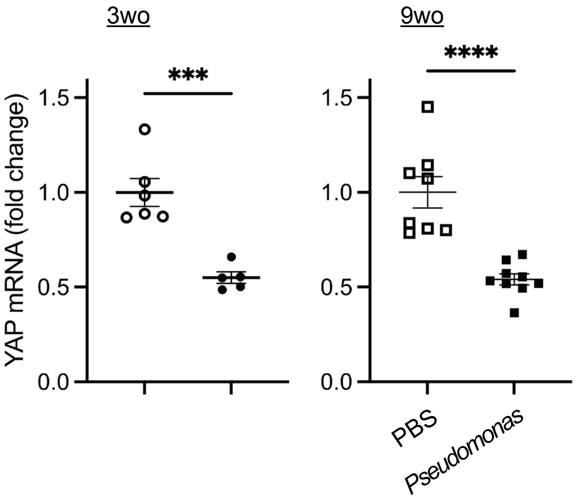

**Supplemental Figure 4:** NF-κB nuclear translocation in weanling mice

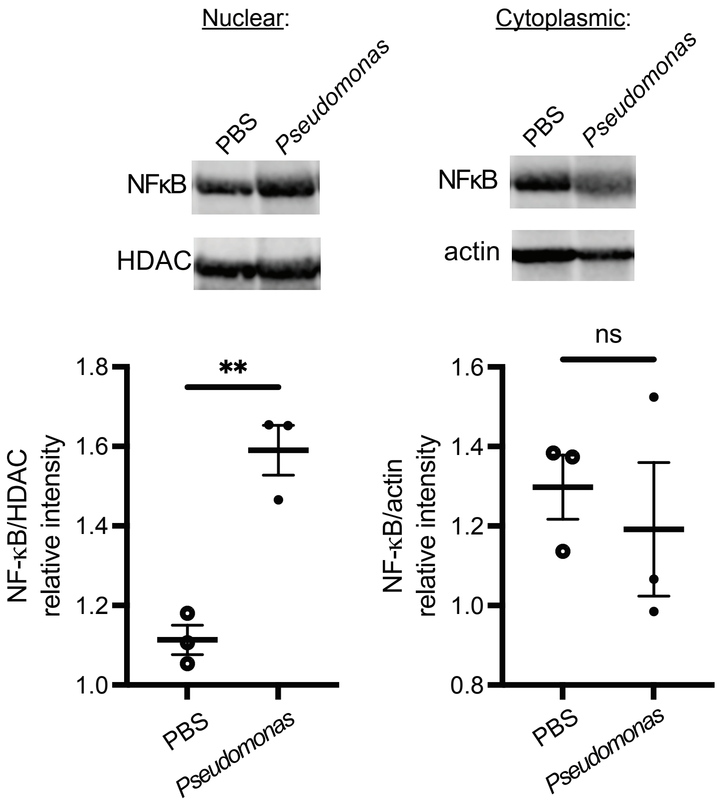
